# A flexible simulation toolkit for designing and evaluating ChIP-sequencing experiments

**DOI:** 10.1101/624486

**Authors:** An Zheng, Michael Lamkin, Yutong Qiu, Kevin Ren, Alon Goren, Melissa Gymrek

**Affiliations:** Department of Computer Science and Engineering, University of California San Diego, La Jolla, CA USA; Department of Bioengineering, University of California San Diego, La Jolla, CA USA; School of Computer Science, Carnegie Mellon University, Pittsburgh, PA USA; Department of Mathematics, Massachusetts Institute of Technology, Cambridge, MA USA; Department of Medicine, University of California San Diego, La Jolla, CA USA

## Abstract

A major challenge in evaluating quantitative ChIP-seq analyses, such as peak calling and differential binding, is a lack of reliable ground truth data. We present Tulip, a toolkit for rapidly simulating ChIP-seq data using statistical models of the experimental steps. Tulip may be used for a range of applications, including power analysis for experimental design, benchmarking of analysis tools, and modeling effects of processes such as replication on ChIP-seq signals.

## Main Text

Chromatin immunoprecipitation followed by next generation sequencing (ChIP-seq) is a widely used technology for genome-wide mapping of the location of DNA-associated proteins, such as transcription factors (TFs), histone modifications (HMs), and chromatin regulators (CRs)^1^. Dozens of methods have been developed for quantitatively analyzing ChIP-seq data, including peak callers (*e.g*., MACS2^2^, HOMER^3^) and differential binding tools (*e.g*. diffbind^4^, DESeq2^5^). A major challenge in training and evaluating these methods as well as interpreting their results is a lack of reliable ground truth data: in most cases, the actual location and strength of binding sites is not known and cannot be reliably measured using orthogonal experimental techniques. Computational analysis of ChIP-seq is further complicated by multiple sources of noise introduced during the experimental process, including inefficiency and/or non-specificity of antibodies, PCR artifacts, and sequencing errors^6,7^. Additionally, similar to RNA-seq, enrichments represent fractions of the total read pool, rather than absolute counts^8^, making it difficult or impossible to determine absolute differences in binding between conditions. Accurate simulation of ChIP-seq data can mitigate this challenge, but existing frameworks^9–11^ are either too cumbersome to use or do not capture important sources of variation present in real data such as pulldown non-specificity or fragment length variability. Here, we present Tulip (Toolkit for simULating IP-sequencing data), a flexible toolkit for rapidly simulating ChIP-seq data based on realistic statistical models of key steps of the ChIP-seq process. We demonstrate the utility of Tulip with two use cases: (1) measuring power to detect binding sites under a range of experimental conditions and (2) analyzing the effects of spike-in normalization controls.

We directly model each major ChIP-seq experimental step (**Figure 1a**; steps 1-4). Below, we describe how these parameters are learned from existing datasets, but these may also be manually set by the user. In <u>*step 1*</u>, DNA is sheared to a target fragment length, typically using mechanical or enzymatic approaches^12^. We model fragment sizes using a gamma distribution (**Figure 1b**) based on empirical observation of size distributions which have long right tails (**Supplementary Figure 2**). For paired-end data, the distribution is learned directly from observed fragment lengths. We additionally developed a novel numerical approach to infer fragment length distributions from single-end data (**Online Methods, Supplementary Figure 2**). In <u>*step 2*</u>, immunoprecipitation enriches for fragments bound to a target epitope (*e.g*., TF, CR, or HM). From an input set of peaks, we model the relative probability of pulldown for bound vs. unbound fragments, which is based on the fraction of the genome in association with the target protein and the antibody specificity (**Figure 1c**). To capture differences in binding across peaks, we additionally model the probability that a specific site is bound according to the strength of the signal at each peak (**Online Methods**). In <u>*step 3*</u>, enriched fragments are amplified using PCR. We model PCR duplicate rates using a geometric distribution, which fits well to observed data (**Figure 1d**). Finally, in <u>*step 4*</u>, libraries of DNA fragments are sequenced. We model substitution and indel errors introduced by sequencing.

**Figure 1:**
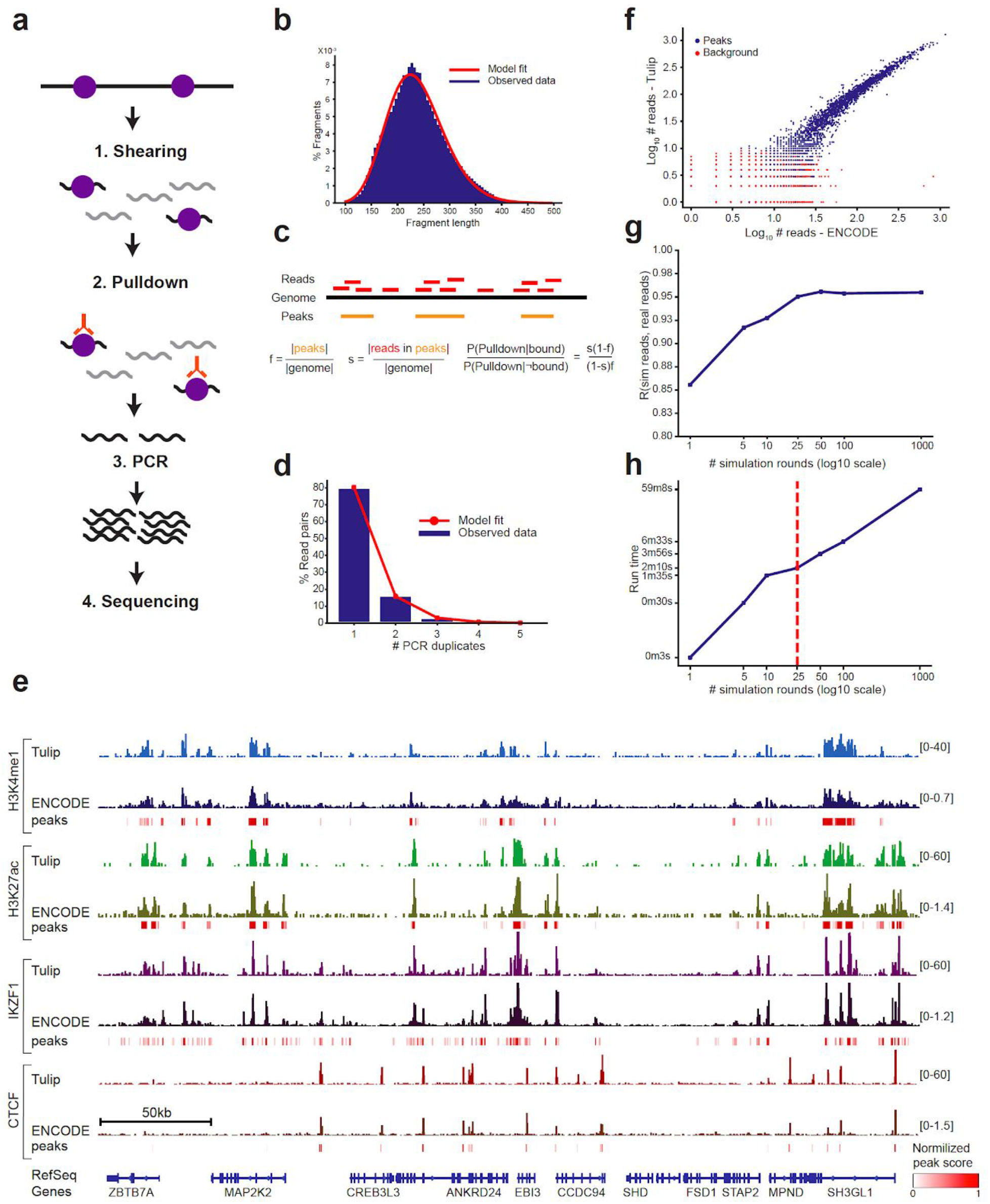
Tulip model and evaluation. **a. Overview of the Tulip model**. Tulip models four steps: shearing, pulldown, PCR, and sequencing. **b. Modeling fragment length as a gamma distribution**. The dark blue histogram shows an example fragment length distribution from real paired end ChIP-seq data. The red line shows the best fit gamma distribution. **c. Schematic of the pulldown model**. Pulldown is modeled using two parameters; *f* (the fraction of the genome bound by the factor) and *s* (the probability that a pulled down fragment is bound. The pulldown model is described in full in the **Online Methods**. **d. Modeling the number of PCR duplicates as a geometric distribution**. The dark blue histograms shows an example of a distribution of the numbers of PCR duplicates in real ChIP-seq data. The red line shows the best fit geometric distribution. **e. Example coverage profiles of real vs. simulated data**. For each HM (H3K27ac and H3K4me1) and TF (CTCF and IKZF1), the bottom track shows peaks identified by ENCODE, the middle track shows coverage profiles based on aligned reads from ENCODE, and the top track shows coverage profiles based on Tulip simulations. Peak scores were normalized to be between 0 and 1 by dividing each peak’s score by 3*the median peak score and thresholding normalized scores at 1 (see color scale in bottom right). Peak score normalization was performed separately for each dataset. We used 25 and 1,000 simulation rounds for HMs and TFs, respectively. Coverage profiles were visualized using the Integrative Genomics Viewer^19^. All tracks were loaded to IGV using the “Normalize Coverage Data” setting. The numbers in brackets to the right of each track show the y-axis scale. ENCODE accessions: H3K27ac (bam=ENCFF097SQI, bed=ENCFF465WTH); H3K4me1 (bam=ENCFF677IFV bed=ENCFF317ATF); IKZF1 (bam=ENCFF216YZE bed=ENCFF795PEX); CTCF (bam=ENCFF598OOE, bed=ENCFF706QLS); **f-h** show data only for H3K27ac **f. Concordance of read counts between simulated vs. real ChIP-seq data**. chr19 was divided into non-overlapping 5kb bins. The scatter plot shows the comparison of read counts per bin for bins overlapping peaks (dark blue) or background regions (red). The x- and y-axes are on a log_10_ scale. The plot shown is for 100 simulation rounds. **g. Read count correlation between real and simulated data as a function of number of simulation rounds**. For each number of rounds, the correlation was computed between read counts in 5kb bins overlapping input peaks. **h. Simulation run time as a function of number of simulation rounds**. Run time is given in CPU-seconds on the y-axis. The red vertical line shows the recommended HM setting using 25 simulation rounds.

Tulip consists of two modules: *simreads* and *learn* (**Supplementary Figure 1**). The *simreads* module takes in ChIP-seq model parameters, including fragment length, pulldown specificity, PCR duplication rate, binding locations and intensities (peaks), plus additional information such as read length and number of reads (**Supplementary Table 1**), and outputs simulated reads. Input parameters can either be set by the user to mimic a future ChIP-seq experiment or learned from existing data using the *learn* module. The user must additionally specify the number of simulation rounds, which denotes the number of times the input reference genome is processed by Tulip. Notably, this number is related to but not directly comparable to the number of experimentally processed cells, since pulldown efficiency is not directly included in our current model (**Online Methods**). We found that using 25-100 simulation rounds works well for HMs and 1,000 rounds works well for TFs (see below).

We evaluated Tulip using ChIP-seq datasets generated by the ENCODE Project^13^ for a variety of TFs and HMs. We learned models for 399 TF and 21 HM datasets from the GM12878 lymphoblastoid cell line (**Supplementary Table 2**). These models can guide design of custom simulations. To evaluate the effect of varying the number of simulation rounds, we simulated reads on chromosome 19 using inferred parameters for H3K27ac in GM12878 over a range of simulation rounds (1-1,000). Run time for chromosome 19 ranged from 3 seconds (1 round) to around 1 CPU-hr (1,000 rounds). Resulting reads were aligned to the hg19 reference genome using BWA-MEM^14^, and duplicates were flagged using Picard (**Online Methods**). Visual inspection of the resulting coverage profiles shows high similarity between real and simulated data (**Figure 1e**). Next, we compared read counts in bins of 5kb and found high correlation between real and simulated data in bins containing at least one peak (Pearson r=0.95, p<10^-200^ for 100 rounds; **Figure 1f**). Bins with no peaks, corresponding to randomly generated background regions, showed no correlation with ENCODE (Pearson r=0.017, p=0.10). Further, correlation with ENCODE data increased as a function of the number of simulation rounds (**Figure 1g**) but plateaued around 100, suggesting little gain in simulating additional rounds compared to the time tradeoff (**Figure 1h**). We repeated this analysis on multiple additional HMs and TFs in GM12878 with similar results (**Figure 1e, Supplementary Figure 3**).

To compare Tulip to the current state-of-the-art, we evaluated ChIPulate^9^, a recently developed method for simulating TF ChIP-seq, using a comparable set of input conditions (**Online Methods**). ChIPulate showed lower correlation with ENCODE for CTCF (Pearson r=0.52 vs. r=0.84 for ChIPulate and Tulip, respectively, across 1kb bins containing at least one peak) and ran for approximately 80 minutes compared to approximately 20 minutes using recommended Tulip settings for TFs (1,000 simulation rounds). Notably, ChIPulate does not simulate background fragments outside of peak regions (**Supplementary Figure 4a**), a key feature of real ChIP-seq datasets related to the antibody specificity, and does not currently perform well on HM data (**Supplementary Figure 4b**).

We next tested the utility of Tulip in two example applications. First, we performed a power analysis to determine the number of reads required to recover a set of peaks based on model parameters learned from ENCODE for a variety of HMs and TFs which encompass diverse genomic binding patterns. For example, H3K27me3 has broad peaks spanning tens to hundreds of kb, whereas H3K4me3 has narrower binding sites located near transcription start sites^15^. For each DNA associated protein, we simulated datasets with numbers of reads varying from 1 million to 100 million. We additionally simulated the same number of WCE reads for each setting. We used MACS2^2^ to call peaks on the resulting datasets after alignment and duplicate marking (**Online Methods**). Power was measured as the fraction of peaks recovered by the simulated datasets. In line with experimental results based on TFs^6^, the power of simulated ChIP-seq datasets for most factors plateaued after 25-50 million reads. Further, our simulations expectedly predict slightly lower power for marks with broader signals (H3K36me3 and H3K27me3). For example, 10 million reads could capture ~80% of input peaks for H3K4me3 but only ~40% of H3K27me3 peaks (**Figure 2a**).

**Figure 2:**
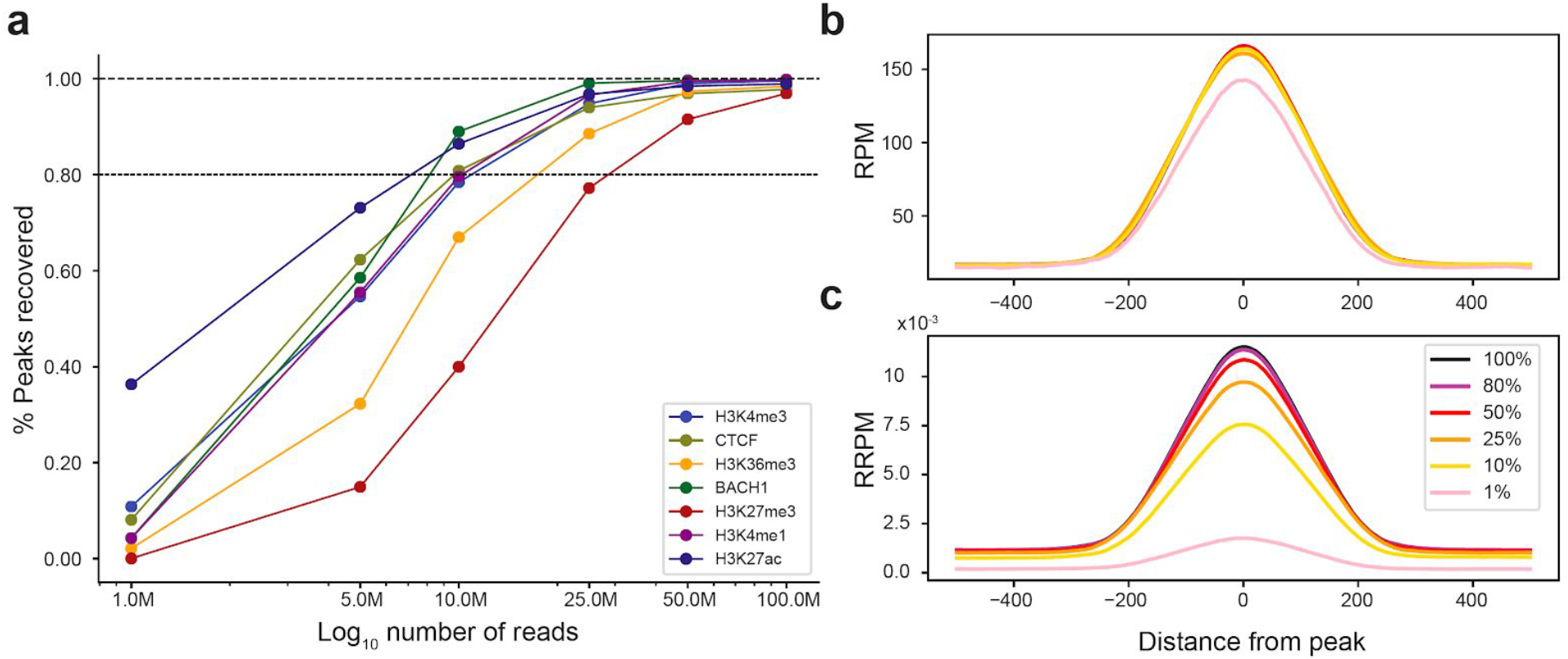
Applications of Tulip for experimental design. **a. Power to detect peaks depends on sequencing depth**. We used Tulip to simulate ChIP-seq datasets using a set of input peaks and model parameters derived from ENCODE datasets. Datasets with different read numbers (1M-100M) were simulated for each TF or HM (colored lines; see legend for details). MACS2 was used to call peaks on the resulting datasets. Power was measured as the fraction of peaks recovered by the simulated datasets. Lower dashed line represents 80% of recovered peaks. The upper dashed line represents 100%. ENCODE accessions used for power analyses are provided in the **Online Methods**. **b,c. Evaluation of spike-in controls**. Tulip was used to generate 6 datasets consisting of peaks with varying binding probabilities (from 100% to 1%) along with a set of “spike in” control regions from *Drosophila* each with constant binding scores of 100%. Coverage profiles centered on the peaks were generated based on two normalization procedures: RPM (upper plot) and reference corrected reads per million (RRPM; lower plot). RPM normalization could not identify the binding changes (except in the most extreme case of a 100-fold reduction (**b**), whereas RRPM correctly distinguished the spectrum of binding levels (**c**).

Next, we used Tulip to evaluate the use of spike-in controls to determine global changes in binding levels of a DNA-associated protein across conditions. Multiple efforts^16–18^ have shown that spiking in a known amount of chromatin from an orthogonal species such as *Drosophila* can be used to more accurately compare absolute binding levels across experiments. To recapitulate this process, we generated a random set of 10,000 TF binding sites from human (chr19) and 20 binding sites from *Drosophila* (chr4). We varied the binding probability of simulated human peaks from 100% down to 1% representing a global reduction in binding affinity and kept the binding probability of *Drosophila* binding sites constant. We simulated ChIP-seq datasets from the spike-in control experiment and aligned reads to a modified reference genome consisting of both species (**Online Methods**). We generated coverage profiles of aligned reads around input peaks based on two normalization procedures: reads per million (RPM) and the reference corrected reads per million (RRPM) metric that employs the spike-in data for normalization^18^. Only RRPM accurately distinguished the spectrum of binding levels (**Figure 2b-c**). This analysis demonstrates Tulip’s utility for exploring *in silico* the effects of spike in and normalization procedures using a ground truth set of binding sites and intensities.

In summary, we present Tulip, an efficient ChIP-seq simulation framework that generates realistic datasets over a flexible range of experimental conditions. Our modular approach allows for future integration of alternative models of different steps, such as modeling effects of GC content or DNA accessibility^7^ on pulldown efficiency. Tulip can generate simulated data for TFs, HMs, and CRs and runs in just seconds to minutes for most applications. We demonstrated the utility of Tulip by measuring power to detect binding sites for TFs and HMs and analyzing the effects of spike-in normalization controls. Tulip may be used for a variety of additional applications including evaluation of computational methods such as peak callers; performing power analyses to determine the number of experimental replicates needed for an analysis; or evaluation of the effects of genetic variation on observed ChIP-seq signals. Overall, we envision our framework will serve as a highly valuable resource for future ChIP-seq analysis efforts.

## Acknowledgements

We thank Drs. Chris Benner and Bing Ren for helpful discussions of the method.

## Author information

### Contributions

A.Z. and M.L. developed statistical models and implemented the Tulip software. Y.Q. conceived statistical models and helped perform benchmarking analyses. K.R. developed a method for estimating fragment length distributions from single end data. A.G. designed the study and wrote the manuscript. M.G. conceived the study and initial statistical models, designed validation experiments, and wrote the manuscript.

### Competing interests

The authors declare no competing financial interests.

## Online Methods

### Tulip model

Tulip models each major step (shearing, pulldown, PCR, and sequencing) of the ChIP-seq protocol (**Figure 1a, steps 1-4**) as a distinct module. It assumes binding sites for the target epitope and their binding scores (probabilities) are known. Model parameters are summarized in **Table 1**.

#### 1. Shearing

In step 1, cross-linked DNA is sheared to a target fragment length, for instance by sonication. Tulip models the length distribution of fragments. We model fragment lengths using a gamma distribution (**Figure 1b**) based on empirical observation of fragment distributions which have long right tails (**Supplementary Figure 2**). In the case of paired-end reads, fragment lengths can be determined trivially from the mapping locations of paired reads. For single-end reads, individual fragment lengths are not directly observed. We outline a novel method below for inferring summary statistics for the length distribution using single-end reads.

##### Inferring fragment lengths from paired-end reads

The observed fragment length (*X_i_*) for each read pair i can be computed based on the mapping coordinates of the two reads. The learn module randomly selects 10,000 read pairs from the input BAM for fitting a gamma distribution. Read pairs are filtered to remove fragments that are unaligned, not properly paired, marked as duplicates or marked as secondary alignments. Read pairs are further filtered to remove fragments with length greater than 3 times the median length of selected fragments.

The mean fragment length is easily computed as 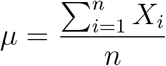, where *n* is the number of fragments remaining after filtering. We then use the method of moments to find maximum likelihood estimates of the gamma distribution shape (*k*) and scale parameters (*θ*):

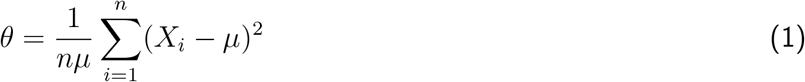

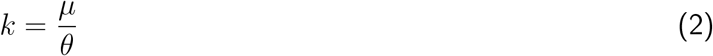

##### Inferring fragment lengths from single-end reads

To estimate the fragment length distribution from single-end reads, we assume the length distribution follows a gamma distribution with mean *μ* and variance *v*, and use reads located inside ChIP-seq peaks (provided as input) to estimate *μ* and *v* which are used to compute *k* and *θ*.

For each peak *peak_i_*, we keep track of two lists, {start}_peak_*i*__ and {end}_peak_*i*__. For each read overlapping peak_*i*_, if the read is on the forward strand we add its start coordinate to {start}_peak_*i*__. If the read is on the reverse strand we add its start coordinate to {end}_peak_*i*__. The center point of this peak is calculated as:

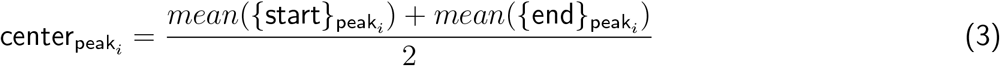

For every peak_*i*_ we offset the coordinates in {start}_peak_*i*__ and {end}_peak_*i*__ by center_peak_*i*__, so that the coordinates of start points and end points are symmetric around zero. We then concatenate lists from each peak to form
{start} and {end}:

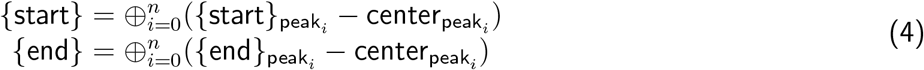

The mean fragment length *μ* can be estimated as:

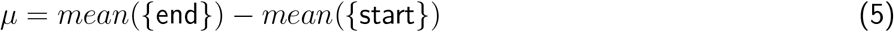

We calculate the probability density functions, cumulative density functions and expected density functions for both {start} and {end}. The expected density function *EDF*(*x*) is defined as the expected deviation of a random element in the list to *x*:

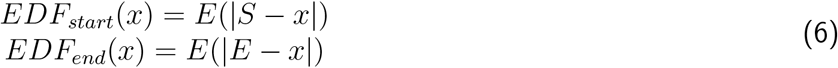

where *S* is a random element in {start} and *E* is a random element in {end}.

After we compute *μ*, we can reduce the density function of the fragment length distribution from *p_μ,v_*(*x*) to *p_v_*(*x*), the probability density function of the fragment length variance. We construct a score function *F*(*v*) as shown below. Intuitively, if we have a correct guess of *v, F*(*v*) should be equal to zero.

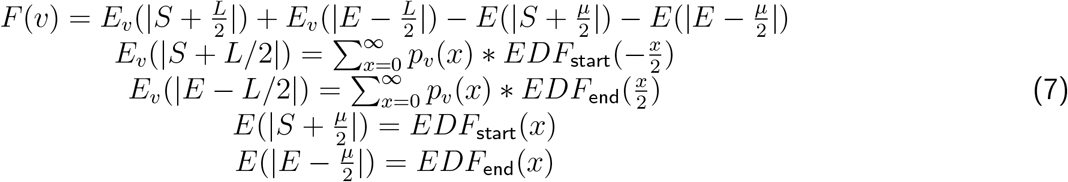

where *L* represents a randomly chosen fragment length.

To find an optimal *v* that minimizes |*F*(*v*)|, we conduct a binary search between 1000 and 10,000.

In practice, we slightly offset the last two items in the score function in **Equation 7** to get the score below, which gives slightly more accurate estimation of v on real data. This may be due to the fact that fragment length distributions are truncated on the left end, with little or no fragments with lengths less than 100bp observed, and thus do not follow a true gamma distribution.

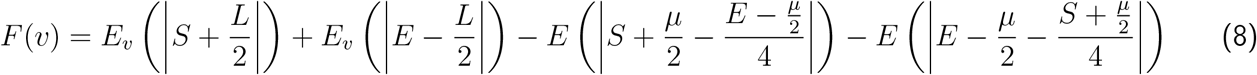

#### Pulldown

In step 2, sheared cross-linked DNA is subject to pulldown, during which an antibody for the protein or modification of interest is used to enrich the pool of fragments for those bound to the epitope recognized by the antibody. This process is imperfect: some bound fragments will not be pulled down, and some unbound fragments will be pulled down. To model this process, we quantify the ratio of the probability of pulling down a bound vs. unbound fragment as described below. This value is specific to each ChIP-seq experiment and depends on the antibody specificity as well as the fraction of the genome bound by the factor of interest.

We use *B_i_* to denote that fragment *i* is bound, 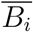 to denote that fragment *i* is unbound, and *D_i_* to denote that fragment *i* is pulled down. For fragment *i*, assuming the probability that a given bound fragment is pulled down is approximately constant over all fragments, the probability of being pulled down can then be written as:

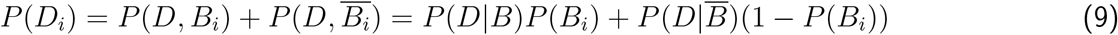

where *P*(*D*|*B*) denotes the probability that a given bound fragment will be pulled down and 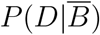 denotes the probability that a given unbound fragment will be pulled down.

For any fragment *i*, we set the probability of it being bound *P*(*B_i_*) based on the scores of peaks it overlaps:

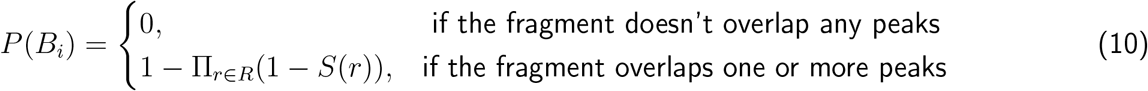

where *R* is the set of all peak regions overlapping fragment *i* and *S*(*r*) is the probability that peak *r* is bound. A method for estimating *S*(*r*) is detailed in the “Tulip implementation” section below.

We compute conditional pulldown probabilities using Bayes’ Rule:

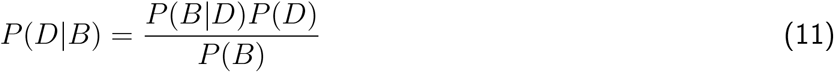

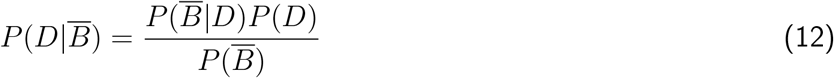

where here *P*(*B*|*D*) and 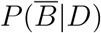 represent averages across all fragments. We do not have a straightforward way to compute *P*(*D*), the average probability that a fragment is pulled down, using only observed ChIP-seq reads since we do not actually observe fragments that are not pulled down. Thus, we do not attempt to compute these conditional probabilities directly. Instead, we take the ratio which cancels *P*(*D*):

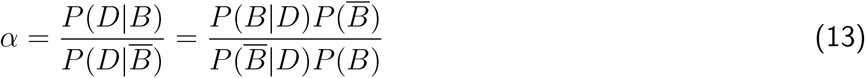

*P*(*B*), or the probability that a fragment is bound on average, is equal to *f*, the fraction of the genome bound by the factor of interest. We can approximate *f* as the sum of the lengths of all peaks *l_k_* weighted by their binding probabilities divided by the total length of the genome *G*:

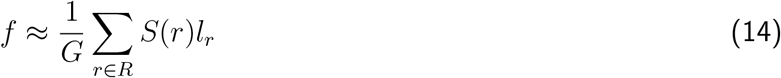

where *G* is the size of the reference genome, *S*(*r*) is the probability peak region *r* is bound as described above, and *l_r_* is the length of peak *r*, assuming no overlap between peaks.

*P*(*B*|*D*), or the probability that a fragment is bound given that it is pulled down, is a measure of the specificity of the antibody. Assuming the majority of reads falling in peaks are truly bound, we can roughly approximate this as the percent of fragments falling within peaks, which we denote as *s*.

Using these two metrics, *f*, and *s* (**Supplementary Table 1**), we can simplify the ratio *α* as:

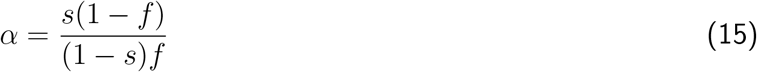

Since we only know the ratio *α*, for simplicity in simulations we set *P*(*D*|*B*) = 1 and 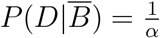. Substituting into Equation 9 above, we compute the probability that fragment *i* is pulled down as:

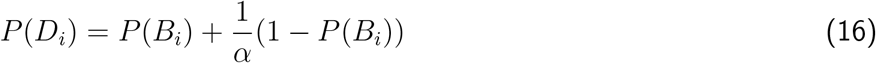

where *P*(*B_i_*) is based on peak scores as defined above.
Note, in reality, *P*(*D*|*B*) is likely to be much less than 1, since the pulldown process is inefficient and many fragments are lost. Further, the number is likely to vary largely across different experiments. Estimating the absolute value of *P*(*D*|*B*) from real datasets is a topic of future work.

#### PCR

In step 3, PCR is used to amplify pulled down fragments before sequencing. Let *n_i_* represent the number of reads (or read pairs) with *i* PCR duplicates (including the original fragment). *n_i_* is modeled using a geometric distribution, where *p* gives the probability that a fragment has no PCR duplicates. The parameter *p* is estimated as 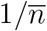, where 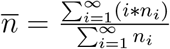.

#### Sequencing

In step 4, amplified fragments are subject to sequencing. Sequences are based on an input reference genome using the coordinates of each fragment. We model the per-base pair substitution rate, insertion rate, and deletion rate (**Supplementary Table 1**).

### Tulip implementation

#### Peak scores

A peak score (*S*(*r*)) is defined for each input peak, where *S*(*r*) gives the probability that a fragment overlapping the region is bound. Note we cannot directly estimate this probability from bulk ChIP-seq data, but assume variability in intensity across peaks is representative of different relative binding probabilities. Based on user input, Tulip computes these binding probabilities either based on peak intensities given in an input BED file or based on read counts from an existing BAM file. If peak intensities are provided, *S*(*r*) is computed as the score of peak *r* divided by the maximum peak intensity. If a BAM file is provided, *S*(*r*) is defined as the number of reads overlapping peak *r* divided by the maximum number of reads overlapping any peak. In both cases, resulting scores *S*(*r*) are between 0 and 1.

By specifiying the option --no-scale, users may directly input binding scores that will be treated directly as probabilities and will not be rescaled. Users may also specify --scale-outliers to remove peak intensities greater than 3 times the median score before rescaling peak intensities to reduce the effect of outlier peaks. In this case all peaks with scores greater than 3 times the median will be set to 1.

#### Learn implementation

The Tulip learn module takes a set of input peaks (BED) and aligned reads (BAM) and learns parameters for (1) fragment length distribution (*k, θ*), (2) pull-down efficiency (*f, s*), and (3) PCR efficiency (*p*) (**Supplementary Table 1**). BAM files must have PCR duplicates marked using a tool such as Picard [1] to enable accurate estimation of PCR parameters. The learn module outputs model parameters to a JSON file that can be used as input to the simreads module.

#### Simulation implementation

The simreads module takes in a set of peaks, model parameters (*e.g*., from the learn module), and experimental parameters (including read length, number of reads) and simulates raw sequencing reads in FASTQ format. The user must additionally specify the number of simulation rounds, which denotes the number of times the input reference genome is processed by Tulip. Notably, this number is related to but not directly comparable to the number of experimentally processed cells, since pulldown efficiency is not directly included in our current model We have found that values of 25-100 simulation rounds work well in most settings for HMs, and 1,000 rounds works well for TFs.

First, each copy of the genome (equivalent to one simulation round) is randomly fragmented based on the specified fragment length gamma distribution. Next, we decide whether to pull down each fragment based on its overlap with input peaks based on *P*(*B_i_*) described above. Then, pulled down fragments are subject to PCR based on the specified PCR rate. Finally, we generate reads from the resulting pool of fragments based on input parameters (using the specified mode read length, number of reads, and mode [single/paired]).

In practice, in each round the genome is processed in bins (default size 100kb) to avoid storing large fragment pools in memory. In a preprocessing step, we determine how many reads to generate from each simulation round based on the target number of output reads. Tulip is parallelized by performing different simulation rounds on separate threads simultaneously.

#### C++ implementation details

Tulip is implemented as an open source C++ project with source code publicly available on Github: https://github.com/gymreklab/Tulip. It uses the open source libraries htslib (www.htslib.org) for parsing BAM files and POSIX threads for multithreading. It supports standard file formats including BAM/CRAM for aligned reads, BED for peak files, JSON for model files, and FASTQ for raw reads. All analyses were performed using Tulip v1.9.

### Inferring parameters of ENCODE datasets

We used the ENCODE Project’s REST API to write a Python script to automatically identify ChIP-sequencing datasets available for the GM12878 cell line. For each replicate of each experiment, we fetched unfiltered BAM files and narrowPeak or broadPeak peak files for transcription factors and histone modifications, respectively (output type=”peaks” or “peaks and background as input for IDR”. We additionally fetched narrowPeak/broadPeak files based on combined replicate information (output type = “optimal idr thresholded peaks”) for use as ground truth peak sets for power analyses.

We used the Picard [1] MarkDuplicates tool (v2.18.11) with default parameters to mark duplicates in each BAM file. For paired end datasets, we called Tulip’s learn module with non-default parameter --paired. For single end datasets, we used non-default parameters -c 7 --thres 100 for transcription factors and -c 7 --thres 5 for histone modifications. For 6 example paired-end datasets, we called learn in both paired end and single end mode to compare inferred fragment length distributions (**Supplementary Figure 2**).

Inferred parameters as well as links to JSON model files that can be used as input to the Tulip simreads module are provided in **Supplementary Table 2**.

### Evaluating Tulip using ENCODE datasets

We performed simulations using an increasing number of simulation rounds (Tulip argument --numcopies 1, 5, 10, 25, 50, 100, or 1000) for two factors (the histone modification H3K27ac and the transcription factor CTCF) with ENCODE data available in the GM12878 cell line. (BAM accession ENCFF097SQI and broadPeak [BED] accession ENCFF465WTH for H3K27ac; BAM accession ENCFF598OOE and narrowPeak [BED] accession ENCFF706QLS for CTCF. For each number of simulations in each factor, we ran Tulip’s simreads tool on hg19 chromosome 19 (--region chr19:1-59128983) using the corresponding learned ENCODE model (see **Supplementary Table 2**) and the same sequencing parameters as the ENCODE reads from chromosome 19 for each dataset (--numreads 269113 --readlen 51 for H3Ka7ac; --numreads 521557 --readlen 36 for CTCF). A similar process was followed for additional datasets displayed in **Figure 1e** (H3K4me1 and IKZF1). We additionally used options -p ENCODE.bed -t bed -c 7, where ENCODE.bed represents the corresponding peak file for each factor with accessions listed above. Simulated reads were aligned to the hg19 reference genome using BWA MEM [2] v0.7.12-r1039 with default parameters and converted to sorted and indexed BAM files using samtools [3] version 1.5. Duplicates were flagged using the Picard [1] MarkDuplicates tool (v2.18.11) with default parameters.

We used the bedtools [4] (v2.27.1) makewindows command to generate a list of non-overlapping windows cross chromosome 19 and the bedtools multicov command to count the number of reads from each BAM file falling in each window. We used bins of size 5kb to evaluate H3K27ac and 1kb to evaluate CTCF, since HMs tend to have much broader peaks than TFs. We used the bedtools intersect command to determine the intersection of each bin with the input peak files. We determined the Pearson correlation between log10 read counts in each bin for the simulated vs. ENCODE dataset after removing bins with 0 counts in each dataset and adding a pseudocount of 1 read to each bin.

Timing experiments were performed in a Linux environment running Centos 7.4.1708 on a server with 28 cores (Intel_®_ Xeon_®_ CPU E5-2660 v4 @ 2.00GHz) and 125 GB RAM using the UNIX “time” command and are based on the “sys” time reported.

### Comparison to ChIPulate

We compared performance ChIPulate [5] vs. Tulip on CTCF (bam=ENCFF406XWF bed=ENCFF833FTF) and H3K27ac (bam=ENCFF385RWJ bed=ENCFF816AHV).

For ChIPulate, we first computed binding energies for each peak required as input. Binding probabilities *S*(*r*) for each peak *r* were computed as described above by scaling read counts from the ENCODE BAM files in each peak to be between 0 and 1. Then we computed binding energies for each region *r* as

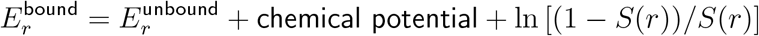

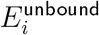 and chemical potential are provided as command line inputs to ChIPulate. We set chemical potential to 0, which provided the best dynamic range across peak scores and highest correlation with real data. Background binding energy was set to the default (1.0).

ChIPulate parameters were set to achieve similar fragment length distributions, PCR rates, number of reads, and read length used for the ENCODE data and Tulip simulations. We used the following settings for running ChIPulate to simulate CTCF based on an input set of peaks identified by ENCODE (accession ENCFF833FTF): p_amp=0.93, p_ext=0.54, --mu-A=0, -d=332, --fragment-length=182, --fragment-jitter=39, --read-length=36. We used the following settings for running ChIPulate to simulate H3K27ac based on an input set of peaks identified by ENCODE (accession ENCFF816AHV): p_amp=0.95, p_ext=0.54, --mu-A=0, -d=101, --fragment-length=211, --fragment-jitter=90, --read-length=51. In both cases we used 1 million cells (-n 1000000) but found results and run time did not vary noticeably when using fewer cells.

To run Tulip on the same dataset, we first learned parameters using the ENCODE BAM and peak files, then ran simreads using the learned model and peak files as input, with parameters: --numcopies 1000 --numreads 521557 --readlen 36 for CTCF and --numcopies 25 --numreads 269113 --readlen 51 for H3K27ac.

Read counts in windows of 1kb and 5kb for CTCF and H3K27ac respectively were compared between simulated and ENCODE datasets using the bedtools multicov command as described above.

### Power analysis

We used Tulip’s simreads tool to generate genome-wide ChIP-seq datasets for five histone modifications (HMs) (H3K27ac, H3K4me3, H3K4me1, H3K27me3, and H3K36me3) and two transcription facetors (TFs) (BACH1 and CTCF) based on model parameters (*f* and *s*) learned from available ENCODE datasets BAM/BED accssions ENCFF385RWJ/ENCFF816AHV, ENCFF398NET/ENCFF795URC, ENCFF809FFP/ENCFF851UKZ, ENCFF677IFV/ENCFF921LKB, ENCFF191SDM/ENCFF479XLN, ENCFF518TTP/ENCFF012JXJ, and ENCFF406XWF/ENCFF833FTF for H3K27ac, H3K4me3, H3K27me3, H3K4me1, H3K36me3, BACH1, and CTCF, respectively. BED accessions are for broadPeak files for the HMs and narrowPeak files for the TFs.

For each HM and TF, we used options -p ENCODE.bed -t bed -c 7 where ENCODE.bed represents the corresponding peak file for each factor with accessions listed above and set --gamma-frag 10,20 --pcr_rate 0.85 --numcopies 100 --readlen 100 to keep these parameters constant across all runs. We simulated datasets using numbers of reads (--numreads) set to 1, 5, 10, 50, or 100 million reads. We additionally simulated a control dataset (whole cell extract; WCE) with the same number of reads but using the options -t wce and no input BED file. As described above, simulated reads were aligned to hg19 using BWA-MEM and duplicates were flagged with Picard. We used MACS2 [6] to call peaks on the resulting duplicate-flagged files using whole cell extract reads as a control and with default options except --broad for histone modifications.

We used the bedtools [4] (v2.27.1) intersect command to determine the percentage of input ENCODE peaks overlapping a peak called by MACS2 from the simulated data, using narrowPeak output for transcription factors and broadPeak output for histone modifications.

### Analysis of spike in controls

We used the bedtools [4] (v2.27.1) random command to generate 10,000 random peaks of size 100 from hg19 chromosome 19 and 20 random peaks of size 100 from Drosophila (dm6) chromosome 4. All peaks were initially given a binding score of 1 (meaning 100% probability of binding).

We then used the Tulip uitlity script chipmunk-spike-in.sh to generate a mock reference genome consisting of the concatenated dm6 and hg19 references and a peak file consisting of both the hg19 and dm6 peaks. We then generated peak files representing different conditions with overall reduced binding of the human sites by setting the scores of all peaks to 0.01, 0.1, 0.25, 0.5, or 0.8, representing reduced binding affinity. For each condition, we used Tulip’s simreads tool with options -p concatenated_peakds.bed -t bed -c 4 -f mock_genome.fa --numcopies 100 --noscale --recomputeF --numreads 1000000 and a model file with f: 15, theta: 20, pcr_rate: 1, and s: 0.4. Note the option --recomputeF is a convenience option to recompute the value of f based on the user input set of peaks.

Reads were aligned to the mock reference genome using BWA MEM. Read densities around peaks were visualized using the annotatePeaks.pl script available from HOMER [7] using options -size 1000 -hist 1. Because exactly 1 million reads were simulated in each case, mean read counts per bin represent reads per million (RPM). To compute reference corrected reads per million (RRPM) based on Orlando, *et al*. [8], we divided each value by the number of reads from the condition mapping to the dm6 genome.

---

**Supplementary Figure 1.**
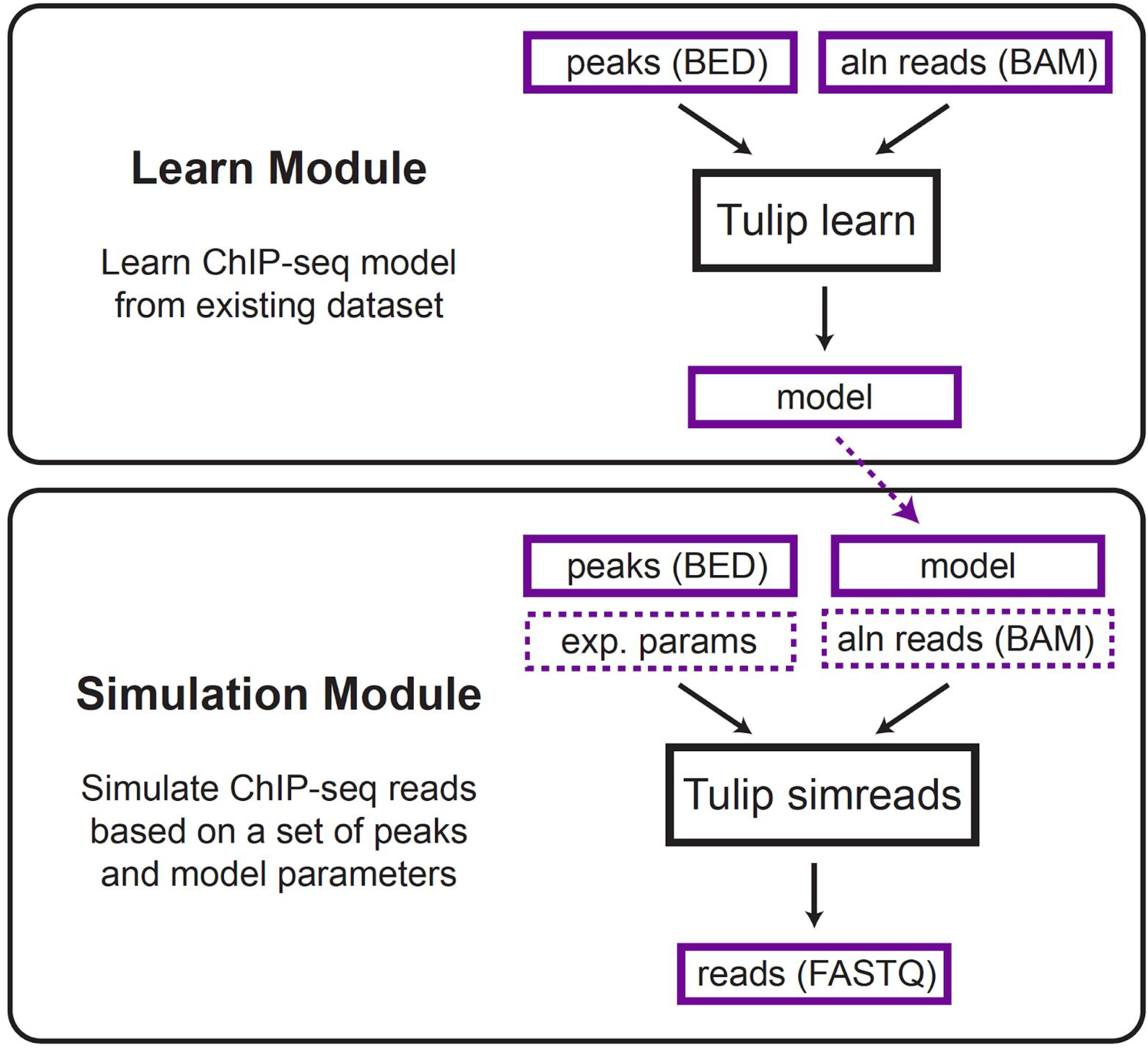
Schematic of Tulip modules. The learn module takes an existing ChIP-seq experiment (aligned reads and peaks) and learns model parameters (see **Supplementary Table 1**). The simulation module takes as input a set of peaks and model parameters, simulates a ChIP-seq experiment, and returns raw reads in FASTQ format. Model parameters input to the simulation module may either be learned from an existing ChIP-seq dataset (dashed arrow) or manually specified to capture planned experimental conditions. Purple borders represent input or output files and black boxes denote Tulip commands. Boxes with solid lines denote required inputs. Boxes with dashed borders denote optional inputs. “Exp. params” denotes experimental parameters including the number of reads, read length, and number of simulation rounds. “Aln reads” denotes aligned reads in BAM format.

**Supplementary Figure 2.**
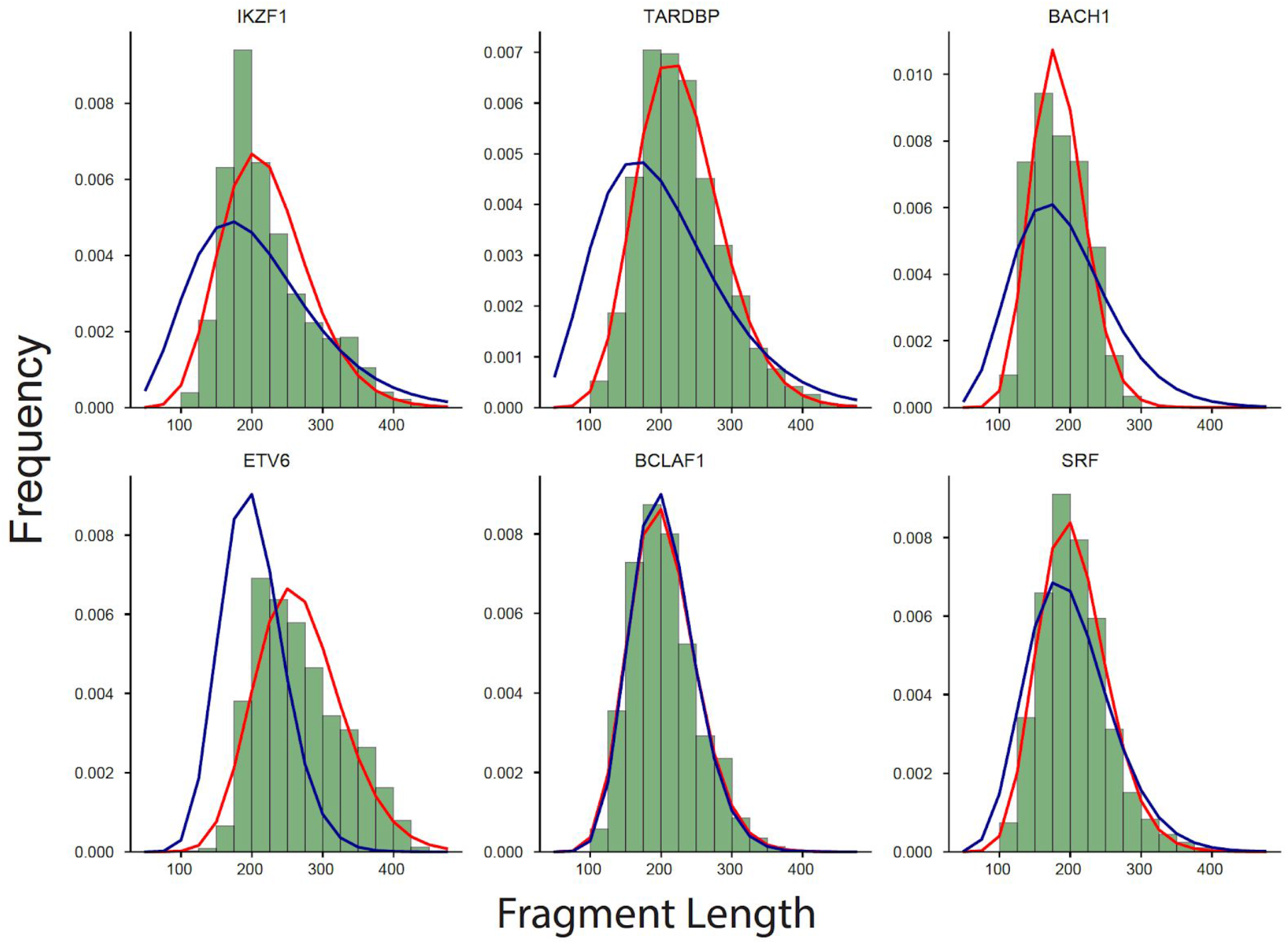
Inferring fragment length distributions from single-end reads. Green bars show a histogram of lengths of 10,000 randomly chosen fragments from GM12878 paired-end ChIP-seq experiments. Red lines give the best fit gamma distribution learned using observed fragment lengths. Blue lines gives the fit inferred ignoring pair information using our novel method for learning fragment length distributions from single end data (**Online Methods**). ENCODE accessions: IKZF1 (bam=ENCFF778KDH bed=ENCFF963HCJ); TARDBP (bam=ENCFF659OTN bed=ENCFF354CAT); BACH1 (bam=ENCFF518TTP bed=ENCFF866OLZ); ETV6 (bam=ENCFF425VPI bed=ENCFF619OWB); BCLAF1 (bam=ENCFF671NSO bed=ENCFF222GJV); SRF (bam=ENCFF387RFR bed=ENCFF500GHH).

**Supplementary Figure 3.**
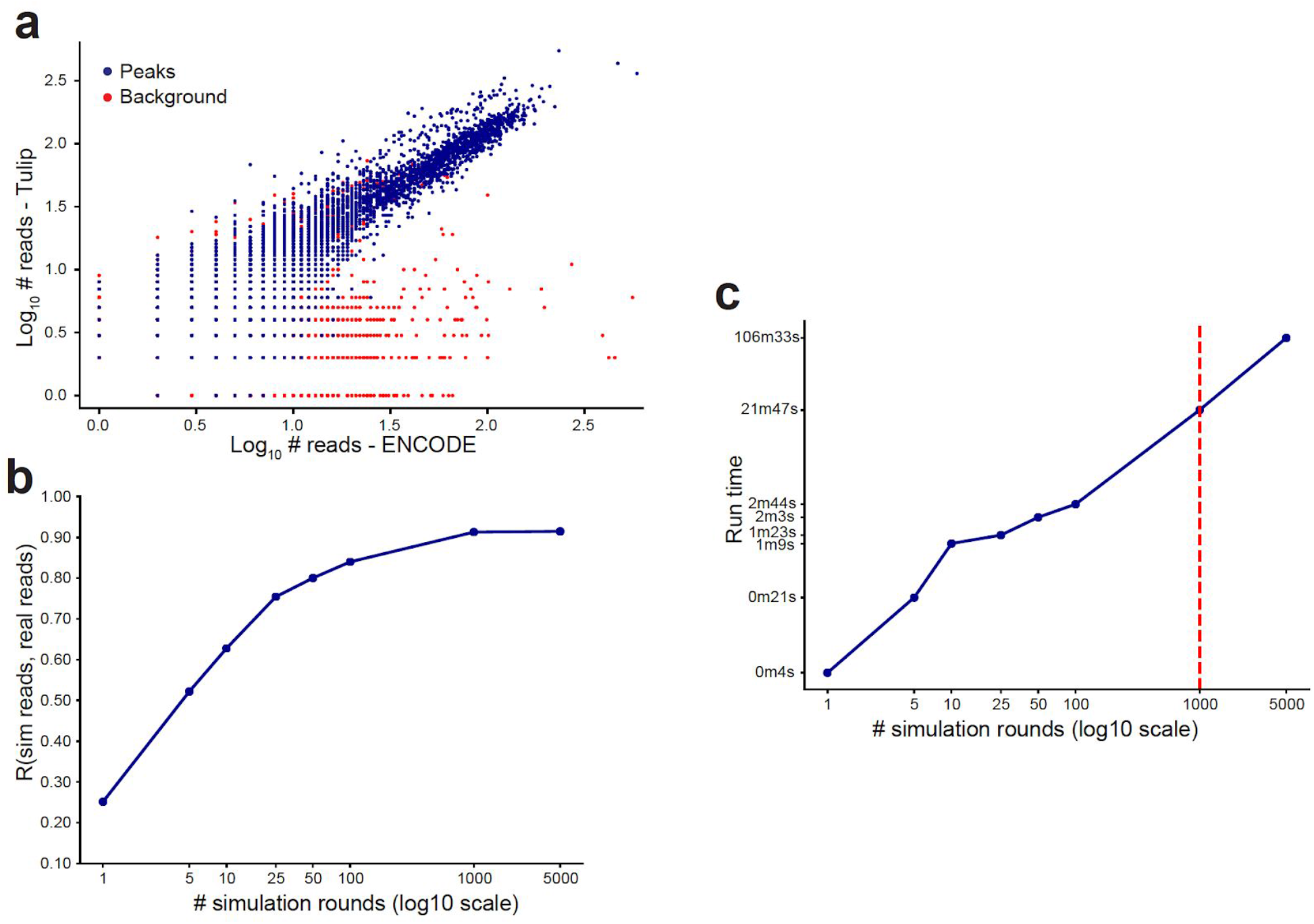
Concordance of real vs. simulated ChIP-seq datasets as a function of number of simulation rounds. **a. Concordance of read counts between simulated vs. real ChIP-seq data.** Paired-end ChIP-seq data for human chromosome 19 was simulated for the transcription factor CTCF using a model learned on an available ENCODE dataset (**Methods**). The chromosome was divided into non-overlapping 1kb bins. The scatter plot shows the comparison of read counts per bin for bins overlapping peaks (dark blue) or background regions (red). The x- and y-axes are on a log_10_ scale. The plot shown is for 1,000 simulation rounds. **b. Read count correlation between real and simulated data as a function of number simulation rounds.** For each number of rounds, the correlation was computed between read counts in 1kb bins overlapping input peaks. **C. Simulation run time as a function of number of simulation rounds.** Run time is given in log_10_ CPU-seconds on the y-axis. The red vertical line shows the recommended TF setting using 1,000 rounds. ENCODE accessions: CTCF (bam=ENCFF598OOE, bed=ENCFF706QLS).

**Supplementary Figure 4.**
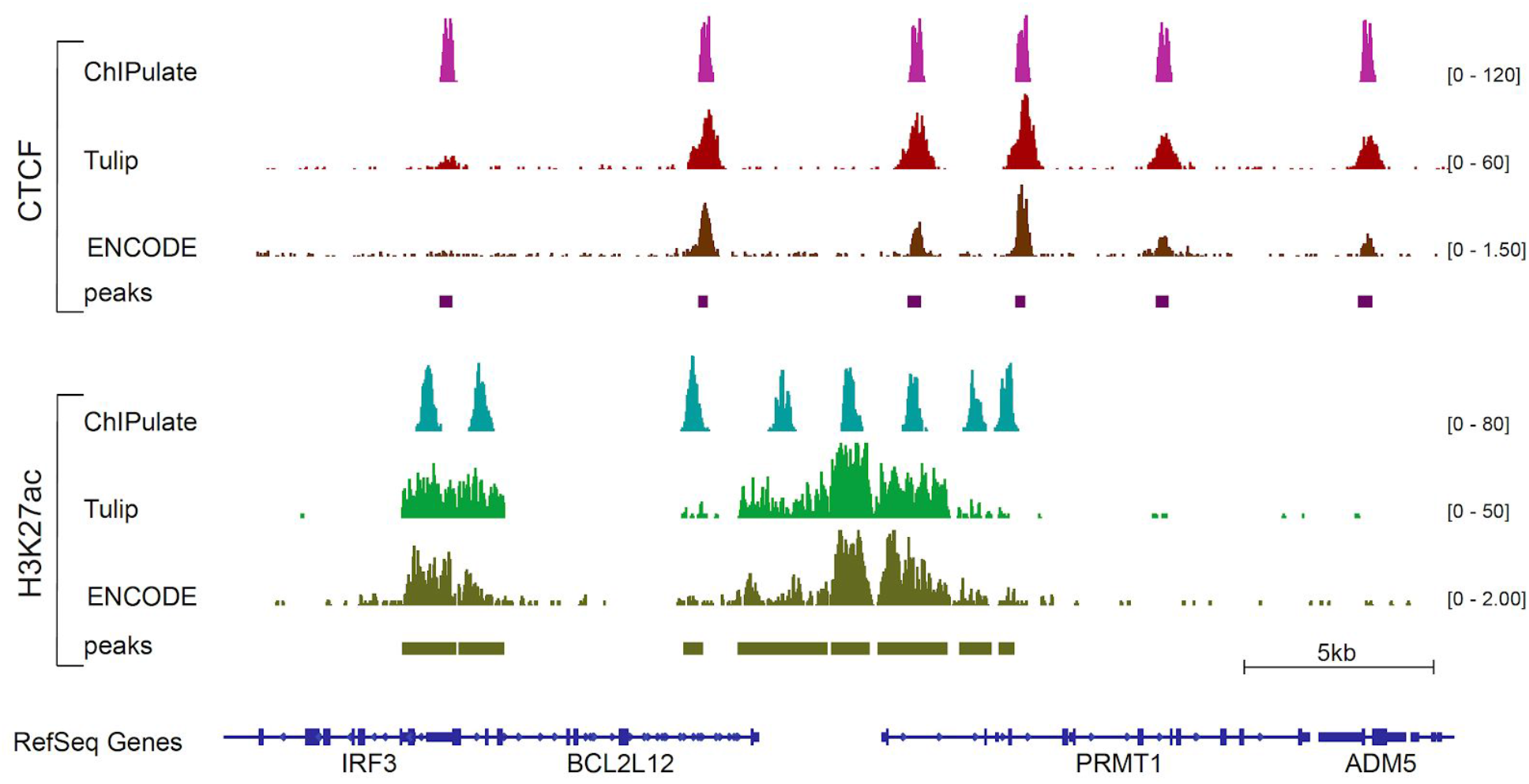
Comparison of coverage profiles for CTCF and H3K27ac from real vs. simulated datasets. For CTCF (top), the bottom bars show ENCODE peaks (accession ENCFF833FTF). The dark brown track shows the ENCODE coverage profile (accession ENCFF406XWF), the dark red track shows the coverage profile obtained using Tulip, and the pink track shows the coverage profile using ChIPulate. For H3K27ac (bottom), the bottom bars show ENCODE peaks (accession ENCFF816AHV). The olive green track shows the ENCODE coverage profile (accession ENCFF385RWJ), the green track shows the coverage profile obtained using Tulip, and the teal track shows the coverage profile obtained using ChIPulate.

**Supplementary Table 1.** Tulip parameters learned or input by users.

| Parameter | ChIP step | simreads argument | Description |
| --- | --- | --- | --- |
| **Parameters learned from real data (learn module)** |  |  |  |
| $k, \theta$ | Shearing | --gamma-frag <float, float> | Parameters of gamma distribution for template lengths |
| $s$ | Pulldown | --spot <float> | SPOT score (percentage of reads falling in peaks) |
| $f$ | Pulldown | --frac <float> | The fraction of the genome bound by the target factor |
| $p$ | PCR | --pcr_rate <float> | The percentage of fragments with no PCR duplicates (geometric distribution parameter). |
| **User-specified parameters** |  |  |  |
| $N$ | Input | --numcopies <int> | The number of simulation rounds, representing the number copies of the genome to process. Higher copy number indicates high number of input cells. |
| $R$ | Sequencing | --numreads <int> | Target number of reads (or read pairs) to simulate. |
| $l$ | Sequencing | --readlen <int> | Length of output reads. |
| - | Sequencing | --paired | Determine whether to output paired vs. single end data. |
| $e, d, i$ | Sequencing | --sub_rate <float><br>--del_rate <float><br>--ins_rate <float> | Sequencing error rates for substitutions, deletions, and insertions. |

**Supplementary Table 2**

See file ZhengEtAl_SuppTable2.xlsx

**Example parameters learned for ENCODE ChIP-seq datasets**.

